# Melanization regulates wound healing by limiting polyploid cell growth in the *Drosophila* epithelium post injury

**DOI:** 10.1101/2025.01.30.635798

**Authors:** Loiselle Gonzalez-Baez, Elizabeth Mortati, Lillie Mitchell, Vicki P. Losick

## Abstract

Wound healing requires a localized response that restricts growth, remodeling, and inflammation to the site of injury. In the fruit fly, *Drosophila melanogaster,* the epithelium heals puncture wounds through cell growth instead of cell division. Epithelial cells on wound margin both fuse and duplicate their genome to generate a multinucleated, polyploid cell essential for tissue repair. Despite the essential role of polyploidy in wound healing, the signals that initiate and regulate the extent of cell growth at the wound site remain poorly understood. One of the first steps in wound healing requires the deposit of melanin at the site of injury, which persists as a melanin scar. The melanin scar forms within hours after a puncture wound and is dependent on the activation of three prophenoloxidase genes (PPO1, PPO2, and PPO3). Using a triple loss of function mutant (*PPO^null^)*, we have uncovered a novel role for melanization in regulating wound healing by limiting polyploid cell growth post injury. Thus, melanization is required for efficient wound closure and its loss leads to an unexpected exacerbation of polyploid cell growth in the surrounding epithelial cells. This occurs, in part, through the early entry of epithelial cells into the endocycle, which may be due to altered gene expression as a result of delayed JNK signaling and other pathways. In conclusion, we have found that polyploid cell growth requires melanization at the injury site to control the extent of cell growth and regulate wound repair.

**SUMMARY:** Puncture wounds in insects are known to lead to production of a pigmented (melanin) deposit that can persist as a scar, remodeling the site of damage. It has long been speculated that the melanin scar is necessary for tissue repair, thus we used the fruit fly as a model to determine how loss of scar formation affects wound healing. In doing so, we discovered that the melanin scar is necessary to restrict cell growth and facilitate efficient wound closure. Thus, revealing a pivotal role for scar formation in coordinating cell growth, which may be conserved in other organisms.

## INTRODUCTION

Polyploid cells, which contain more than the diploid copy of a cell’s genome, have been identified in many tissues across the tree of life (Bailey et al., 2021; Fox et al., 2020). Stress, such as injury, can promote polyploidization which is required for animal tissues to heal and compensate for cell loss. This is the case for several tissues in *Drosophila melanogaster* including the epithelial cells in the abdomen, pyloric cells in the hindgut, and follicle cells in the ovary (Cohen et al., 2018; Losick et al., 2013; Tamori & Deng, 2013). Polyploid cells also contribute to tissue repair in zebrafish epicardium as well as mouse liver and kidney (Cao et al., 2017; De Chiara et al., 2022; Zhang et al., 2018). The polyploid cells arise by whole genome duplication in most injury responses studied to date, except the *Drosophila* epithelium, for which epithelial cells also undergo cell fusion yielding a multinucleated, polyploid cell (Galko & Krasnow, 2004; Losick et al., 2013; White et al., 2023).

Epithelial cells in the adult fly epithelium are post-mitotic and needle puncture wounds were found to heal through endoreplication and cell fusion in a process termed wound-induced polyploidization (WIP) (Losick et al., 2013 and 2016). Epithelial cell fusion is necessary to speed wound healing, whereas endoreplication serves to restore tissue mass. In addition, endoreplication enables healing in the presence of genotoxic stress as well as a means to restore organ mechanics post injury (Grendler et al., 2019; Losick & Duhaime, 2021). What remains poorly understood, however, are the injury signals that initiate polyploidization as well as regulate the extent of cell growth surrounding the wound site. Previous studies found that cell growth is dependent on the conserved signal transduction pathway Hippo, which initiates endoreplication in both *Drosophila* and the mouse kidney via its co-transcriptional activator Yki and Yap, respectively (Besen-McNally et al., 2021; De Chiara et al., 2022; Losick et al., 2013 and 2016). The JNK pathway via its transcription factor Jun, on the other hand, is required to restrict endoreplication (Huang et al., 2024; Losick et al., 2016). Both the Hippo and JNK signal transduction pathways are regulated by extracellular cues, including remodeling of the extracellular matrix (Yu & Guan, 2013). Thus, a plausible hypothesis is to test if scar formation at the site of injury regulates WIP.

Needle puncture wounds lead to deposits of melanin within 1 hour which serves as both a structural scar to repair the insect’s cuticle as well as part of the immune response in defense against infection (Binggeli et al., 2014; Dudzic et al., 2015; Losick et al., 2013; Tang, 2009). Melanin is a pigment whose biosynthesis depends on prophenoloxidase (PPO) genes for which the *Drosophila* genome encodes three: PPO1, PPO2, and PPO3 (Binggeli et al., 2014; Dudzic et al., 2015). PPO1 and PPO2 are expressed in specific hemocytes known as the crystal cells, whereas PPO3 is expressed in lamellocytes. Cell lysis or rupture results in release of the prophenoloxidase proteins, which are then cleaved by extracellular serine proteases upon injury or infection (Dudzic et al., 2015 and 2019; Troha & Buchon, 2019).

Despite melanin’s well characterized role in the immune system, melanin has also been implicated in wound healing by forming a scar at the wound site (Cerenius et al. 2008; Tang et al. 2009). However, its role in regulating epithelial repair and cell growth via polyploidization has remained unexplored. Using a *PPO^null^* strain that harbors loss of function mutations in all three *PPO* genes, we show that loss of melanization leads to a hyper-polyploid response, which includes both exacerbated endocycling and cell fusion post injury. This misregulation of polyploidization appears to be due to altered JNK signaling leading to a defect in wound closure. Thus, we conclude that melanization coordinates signaling pathways that are essential to regulate the extent of polyploidization post injury.

## MATERIALS AND METHODS

### *Drosophila* husbandry and strains

The *Drosophila melanogaster* strains were raised on corn syrup, soy flour-based fly food (Archon Scientific) at 25°C with 60% humidity and 12:12 hour light and dark cycle. *Drosophila* strains for this study were from Bloomington Drosophila Stock Center and Vienna Drosophila Resource Center as noted in the Table S1. The PPO mutant strains (*PPO1^null^*, *PPO2^null^*, and triple *PPO^null^*) were generously provided by Bruno Lemaitre (Binggeli et al., 2014; Dudzic et al. 2015).

### Injury, dissection, immunostaining, and imaging

Adult flies were aged until 3-6 days old and then wounded and dissected as previously described (Bailey et al., 2020). Briefly, dissected abdomens were fixed with 4% paraformaldehyde for 30 minutes, permeabilized in 1 x PBS with 0.3% Triton X-100 and 0.3% BSA, and then stained with primary and secondary antibodies as noted in Table S2. Stained abdomens were mounted in Vectashield (Vector Labs, H-1000) on a glass coverslip and slide, with the inner tissue facing out. Tissue samples were imaged on a Zeiss Axiovert with ApoTome using a 20x or a 40x dry objective. Full Z-stack images were taken at 0.5μm per slice and flattened into a SUM or MAX of stacks projections using FIJI/ImageJ (SCR_002285) imaging software for all analysis.

### Epithelial wound closure assay

Flies expressing a membrane-bound UAS-mCD8RFP under epi-Gal4 control were injured, dissected at the indicated time points, fixed, and stained with antibodies to RFP and FasIII (Table S2). All samples were imaged under the same conditions and settings. Using FIJI, the SUM projection of the RFP channel was used to outline and measure the open area.

### Ploidy Quantification

Abdomens were stained with antibodies to Grh and DAPI dye to identify and measure epithelial nuclear ploidy as previously reported (Bailey et al. 2020). The uninjured control epithelial nuclei were used as an internal control as ploidy has been previously measured to be diploid (2C) (Losick et al., 2013 and 2016). All samples were imaged under the same conditions and settings. Using FIJI, the Grh and DAPI channels were overlaid, and ROIs were drawn around each nucleus that was positive for both the epithelial-specific nuclear marker Grh and DAPI. ROIs were transferred to the corresponding SUM projection of the Z stack of DAPI. The DAPI intensity and nuclear area were measured for each ROI, except for nuclei that overlapped with non-epithelial nuclei, which were excluded from analysis. The ploidy was calculated by normalizing the DAPI intensity of the average value of the 2C uninjured epithelial nuclei for at least three abdomens per condition. The normalized ploidy values were binned into the indicated color-coded groups: 2C (0.5-3.0C), 4C (3.01C-6.0C), 8C (6.01-12.0C), and 16C (12.01C-24C). Ploidy for uninjured samples was measured within a 300 x 175 μm^2^ area while ploidy for injured samples was measured within a 332.80 x 332.80 μm^2^.

### Image analysis

Using FIJI, the syncytium size was quantified by outlining the wound site syncytium using the FasIII cell junction marker as a guide. The number of Grh^+^ epithelial nuclei within the syncytium were quantified using the cell counter tool. For PCNA-EGFP, a 250 x 250μm^2^ ROI box was placed with the wound at the center and overlaid on the SUM of stacks projection of the GFP channel Z-stack. The GFP channel was used to count the number of EGFP-PCNA positive nuclei. For TRE-dsRED, a 330 x 330μm^2^ ROI box was placed with the wound at the center and overlaid on the MAX of stacks projection of the RFP channel Z-stack. Using the cell counter tool in FIJI, nuclei positive for TRE-dsRED expression were counted within the indicated area. Lastly, for MMP1, a 300 x 300μm^2^ ROI box was placed with the wound at the center and overlaid on the SUM of stacks projection of the MMP1 channel Z-stack. The integrated density and area were measured. The MAX of stacks projection was used to outline and measure the MMP1^+^ area.

### Replicates and Statistical Analysis

All experiments were performed in triplicate with at least three biological replicates (fruit flies) per experiment resulting in total n of 10 flies analyzed unless otherwise noted.

Raw values from each graph are available in the Source data 1 - 7 files. Excel was used for basic calculations (ex. ploidy, syncytium size) and statistical analysis was performed using GraphPad Prism (i.e., ANOVA with Tukey’s multiple comparisons test and Student t-test with Welch’s correction) as indicated in the Figure Legend.

## RESULTS

### Melanization is required for wound closure

Melanization occurs within hours at the site of injury and has been speculated to be required for wound healing, but never definitively shown (Cerenius et al., 2008; Tang, 2009). Thus, our first task was to determine if the *Drosophila* epithelium can heal when melanization is inhibited. To do so, we acquired a triple prophenoloxidase mutant strain that has loss of function mutations in all three prophenoloxidase genes (PPO1, PPO2, and PPO3, referred to here as *PPO^null^*) (Dudzic et al., 2015). The *PPO^null^* strain does not produce melanin at the wound site, whereas the wildtype (*w^1118^*) strain generates a robust melanin deposit by 1 hour post injury (hpi) or as shown at 3 days post injury (dpi) (Figure 1a). The epithelial membrane was then visualized by expressing an epithelial specific membrane-tethered RFP using the Gal4/UAS system (Bailey et al., 2020; Phelps & Brand, 1998). The control (*w^1118^; epi-Gal4>UAS-mCD8-RFP*) and *PPO^null^* (*PPO^null^*; *epi-Gal4>UAS-mCD8-RFP*) strains were generated by standard mating schemes in *Drosophila* and assayed for a continuous epithelium. Injury of all fly strains led to a comparable mean wound size of 3,000- 4,000μm^2^ at 1 hpi and consistent with our previous reports, a continuous epithelium was restored in the wildtype strain by 3 dpi (Figure 1b-1e) (Besen-McNally et al., 2021; Losick et al. 2013). Likewise, inhibition of WIP by the simultaneous inhibition of the endocycle (*E2F1^RNAi^*) and cell fusion *(Rac^DN^*), henceforth referred to as a *WIP^null^,* (epi-Gal4> UAS-*E2F1^RNAi^*;UAS-*Rac^DN^*) did not inhibit scar formation (Figure 1a), but resulted in open wounds at 3 dpi (Figure 1c and 1e). As a result, we found that wound healing was significantly impaired in the *PPO^null^* flies, yielding an open area comparable to or larger than the *WIP^null^* strain. Thus, melanization appears to be required for wound healing in *Drosophila* (Figure 1c and 1f).

**Figure 1.**
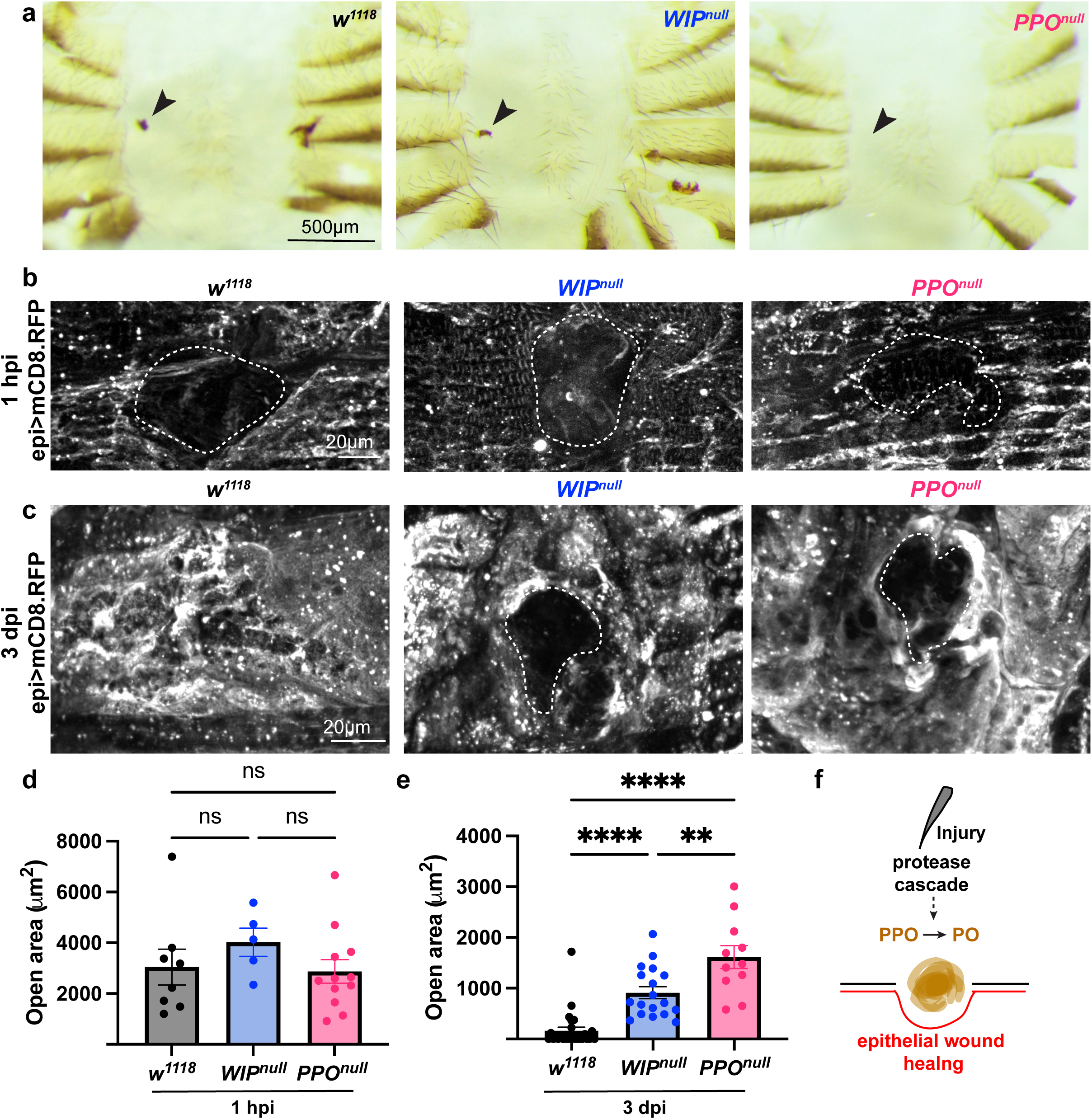
**Melanization is required for wound closure**. (a) Representative brightfield images of dissected abdomens 3 days post injury (dpi) demonstrating that a melanin scar fails to form when all three prophenoloxidase genes are knocked out. (b, c) Representative immunofluorescent images of epithelium in wildtype (*w^1118^*), *WIP^null^,* and *PPO^null^* at (b) 1 hpi and (c) 3 dpi. Epithelial membrane (mCD8.RFP) and open area (outlined, white dashed line) are indicated. (d) Quantification showing that initial wound size is not significantly different in all three strains. (e) Quantification showing that wounds fail to close by 3 dpi when prophenoloxidase is knocked out. (f) Schematic showing phenoloxidase is induced post-injury and is required for epithelial wound healing. Data represent the mean±s.e.m. \*\**p*<0.001; \*\*\**p*<0.0001; ns, not significant (one-way ANOVA with Tukey’s multiple comparisons test). See Source data 1.

### Melanization limits epithelial syncytial size post injury

Our previous studies on wound healing in adult fly epithelium demonstrated that polyploidization is required for tissue repair, thus we hypothesized that melanization was required to stimulate polyploidy either via cell fusion or the endocycle. The epithelium was immunostained with antibodies to epithelial specific proteins, including FasIII (a septate junction protein to visualize cell junctions) and Grainyhead (Grh, a transcription factor that localizes to nucleus of epithelial cells) (Bailey et al., 2020). The wildtype epithelium is mononucleated and loss of melanization in the *PPO^null^* strain did not appear to impact epithelial organization pre-injury (Figure 2a and 2b). However, we found that epithelial cells in *PPO^null^* mutants fused significantly more post injury compared to wildtype (*w^1118^*) flies (Figure 2c and 2d), as indicated by the significant increase in syncytium size and number of nuclei per syncytium (Figure 2e-2g). At first it appeared there were fewer epithelial nuclei within the syncytium in *PPO^null^* flies, but instead this was due to significant down regulation of Grh in epithelial nuclei within the syncytium (Figure 2d’ and 2h). Overall, we observed a surprising exacerbation of the epithelial syncytia size post injury suggesting that melanization is required to limit epithelial cell fusion post injury.

**Figure 2.**
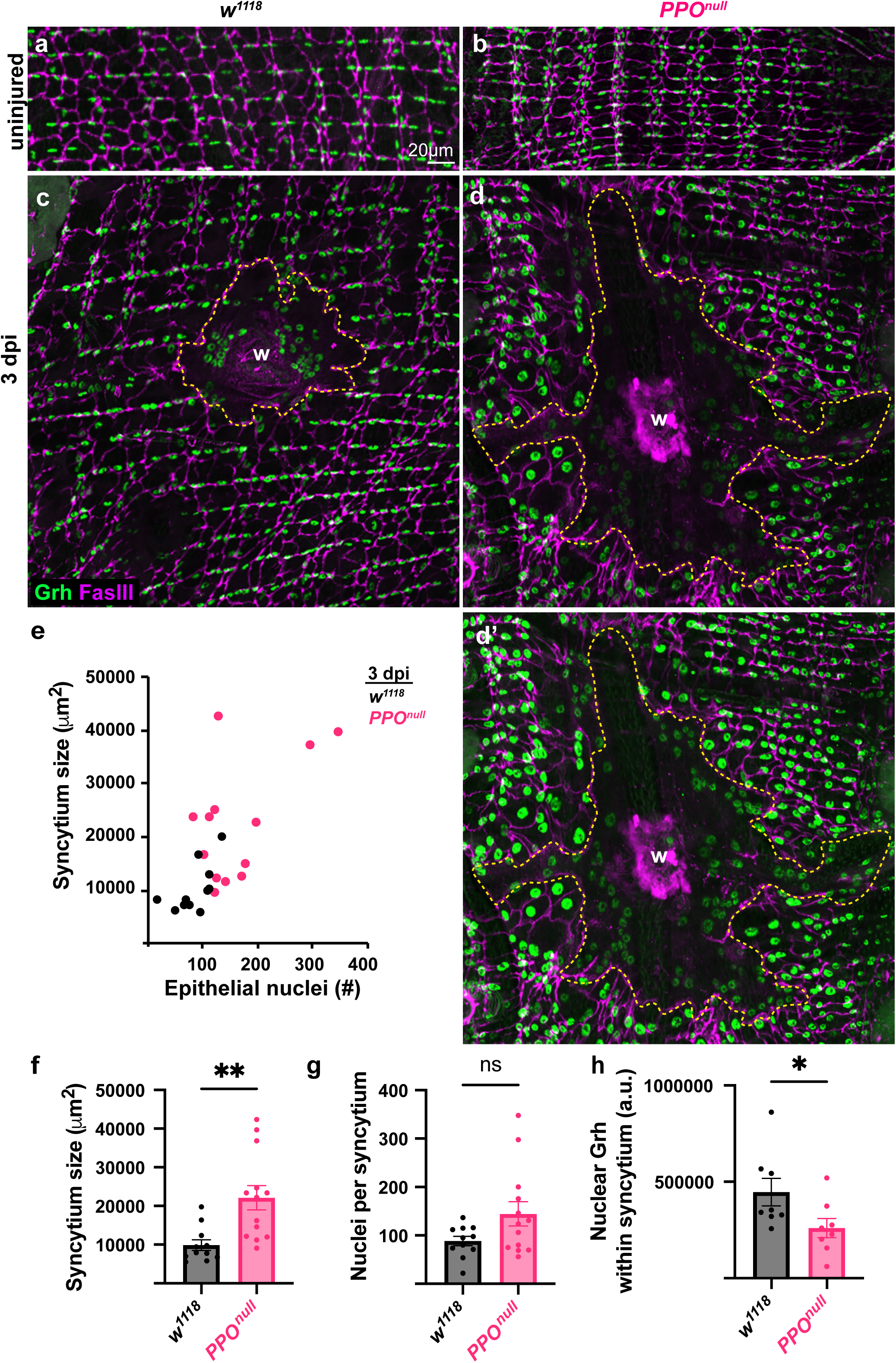
**Melanization limits epithelial syncytial size post injury**. Representative immunofluorescence images of epithelium in uninjured (a) *w^1118^* and (b) *PPO^null^* as well as 3 dpi (c) *w^1118^* and (d) *PPO^null^.* (d’) Brightened 3 dpi *PPO^null^* image to observe epithelial nuclei within the syncytium. Septate junctions (FasIII, magenta), epithelial nuclei (Grh, green), multinucleated cell (outlined, yellow dashed line), and wound site (w) are indicated. (e and f) Quantification showing a larger syncytium size forms post injury when prophenoloxidase genes are knocked out. (g) Quantification of number of nuclei per syncytium. (h) Quantification of nuclear Grh showing it is downregulated in nuclei within the syncytium in the *PPO^null^* fly strain. Data represent the mean±s.e.m. \**p*<0.05 (Student’s t-test). See Source data 2.

To determine if knockout of a single *PPO* gene is sufficient to induce a hyper wound-induced cell fusion response, we measured the syncytium size in individual *PPO* loss of function mutations, referred to here as *PPO1^null^* and *PPO2^null^* respectively (Binggeli et al., 2014). PPO3 is expressed in larval lamellocytes, cells which have not been observed in adult flies, hence we did not investigate the PPO3 gene further.

Knockout of either *PPO1* or *PPO2* did not affect wound size, as both strains had similar sized melanin scars compared to the wildtype (*w^1118^*) (Figure S1a and S1d). Genetic loss of individual PPO genes did not affect epithelial development nor WIP as a similar sized syncytia was generated in epithelium at 4 dpi (Figure S1b-S1f). There was no significant difference in syncytium size nor the number of nuclei between wildtype and *PPO1^null^* or *PPO2^null^* mutants. Thus, we concluded that enhanced polyploidization in the *PPO^null^* flies is more likely due to the loss of melanization, itself, rather than the individual activity of the *PPO* genes.

### Melanization limits epithelial ploidy post injury

The most notable phenotype in the *PPO^null^* flies was a dramatic enlargement of the epithelial nuclei in terms of their size surrounding the giant syncytium at the wound site (Figure 2d). Hence, we investigated if genetic loss of melanization led to exacerbated endoreplication by using our previously reported semi-automated nuclear ploidy analysis (Bailey et al., 2020; Losick et al., 2016). To do so, a region of interest (ROI) was drawn around each nucleus within the epithelium (Figure 3a). The ROIs were then overlaid onto the DAPI image and any overlapping nuclei removed from further analysis. This method allows a majority of epithelial nuclear ploidy to be determined. A heat map of the epithelial nuclear ploidy for each sample was created to analyze the spatial distribution of the nuclei based on ploidy. Pre-injury, about 94% of the epithelial nuclei were diploid in both wildtype and *PPO^null^* flies (Figure 3b, 3c, and 3f). The total number of epithelial nuclei in the *PPO^null^* strain appeared to be enhanced compared to the wildtype, although quantification did not reveal a significant difference (Figure 3h, *P*-value= 0.088). After injury though, ∼22% of the epithelial nuclei in wildtype flies doubled their nuclear ploidy to tetraploid (∼4C) around the wound site (w) (Figure 3d, 3f and 3g). Whereas loss of melanization resulted in a significant increase in polyploid nuclei, with about 42% of nuclei becoming polyploid (≥3C) at 3 dpi (Figure 3e- 3g). These nuclei reached higher ploidy values up to 8C or greater (Figure 3g). In conclusion, these results demonstrate that melanization is required to limit epithelial ploidy post injury as both nuclear ploidy and the number of polyploid nuclei per tissue is exacerbated by the genetic loss of the prophenoloxidase genes in *Drosophila*.

**Figure 3.**
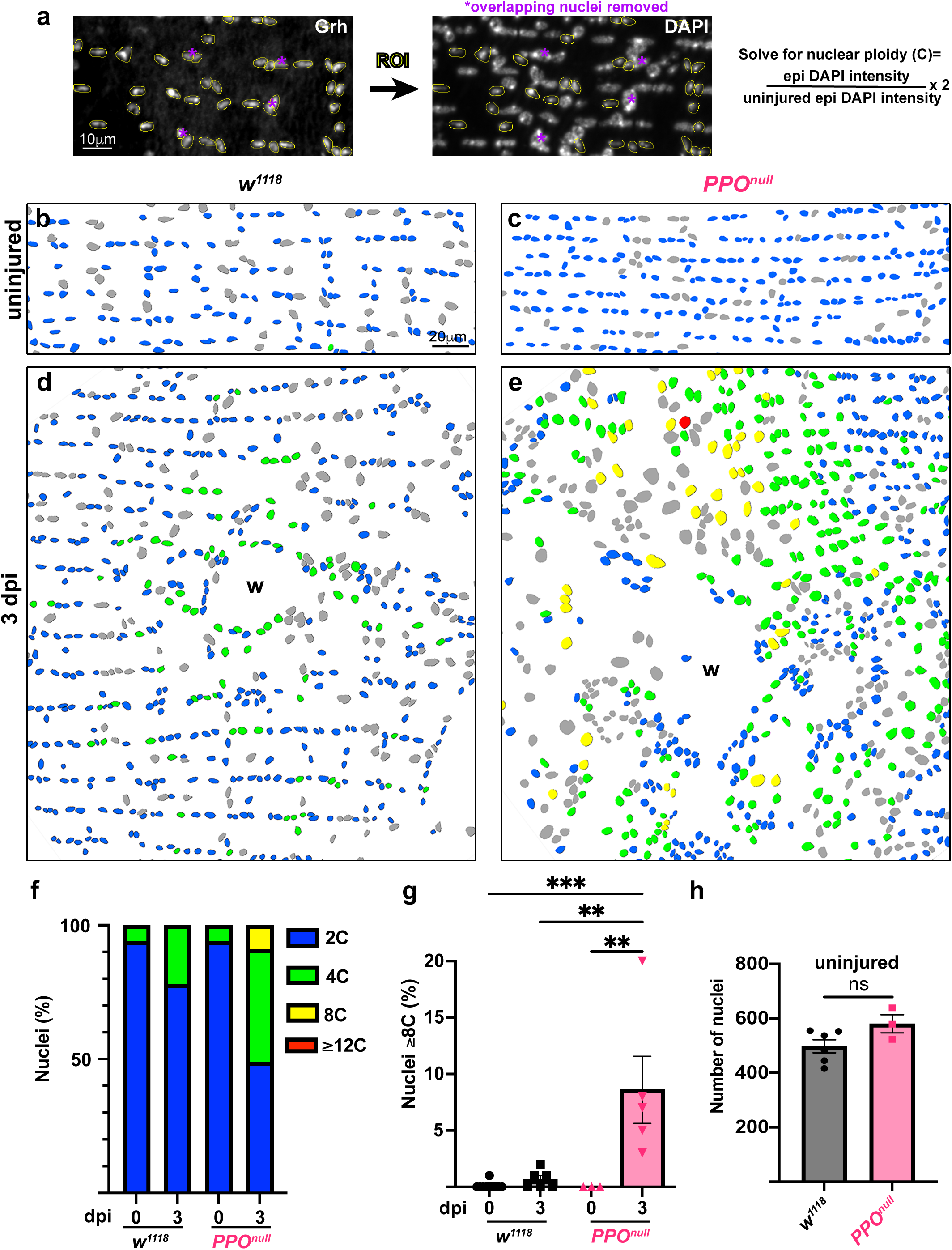
**Melanization limits epithelial nuclear ploidy post injury**. (a) Schematic showing how nuclear ploidy was measured by identifying epithelial nuclei and transferring ROI to the DAPI image. Nuclei overlapping with non-epithelial nuclei were not determined (*ND). Representative heat map images of uninjured (b) *w^1118^* and (c) *PPO^null^* as well as 3 dpi (d) *w^1118^* and (e) *PPO^null^*. Nuclei are colored coded based on ploidy with ND nuclei in gray with wound site (w) labeled. (f) Quantification of percent of epithelial nuclei in the indicated color-coded ploidy ranges. (g) Quantification of percent of nuclei ≥ 8C at 3 dpi. (h) Quantification showing loss of PPO genes does not significantly affect nuclei number pre-injury. Data represent the mean±s.e.m. \*\**p*<0.001; \*\*\**p*<0.0001; ns, not significant (one-way ANOVA with Tukey’s multiple comparisons test). See Source data 3.

### Melanization is required to limit the extent of cell cycle activity

Now that we observed an exacerbated nuclear ploidy response to injury in the absence of melanization, we hypothesized this was due to a difference in cell cycle activity. To determine if the loss of melanization affected cell cycle entry, we generated a recombinant *PPO^null^* strain with the S-phase reporter, EGFP-PCNA (Blythe & Wieschaus, 2016). As expected PCNA was not expressed in epithelial cells pre-injury, indicating that the uninjured epithelium is quiescent same as the wildtype strain (Figure 4a-4c). Our prior studies showed that epithelial cells enter S-phase by 24 hpi and as expected we observed ∼150 PCNA^+^ nuclei in the wildtype flies at this time point (Losick et al. 2013). In the absence of melanization, however, S-phase entry was expedited, with ∼150 nuclei becoming PCNA^+^ by 12 hpi (Figure 4b and 4c). By 24 hpi, ∼500 nuclei were PCNA^+^ in *PPO^null^* flies, indicating an exacerbated response similar to our nuclear ploidy analysis findings (Figure 3, 4b, and 4c). To determine if *PPO^null^* mutants increased nuclear ploidy post injury through an endocycle, which bypasses M phase, or through failed karyokinesis that truncates M phase, we stained for the M-phase marker, Cyclin B (CycB). However, we did not detect any CycB expression in either the wildtype or *PPO^null^* flies post injury. Overexpression of CycB in the fly epithelium (epi-Gal4, UAS- CycB) validated that antibody could detect CycB, which was robustly expressed (Figure 4d). Taken together, this data suggests that melanization regulates the timing and extent of S-phase entry to coordinate and control the extent of polyploidization post injury.

**Figure 4.**
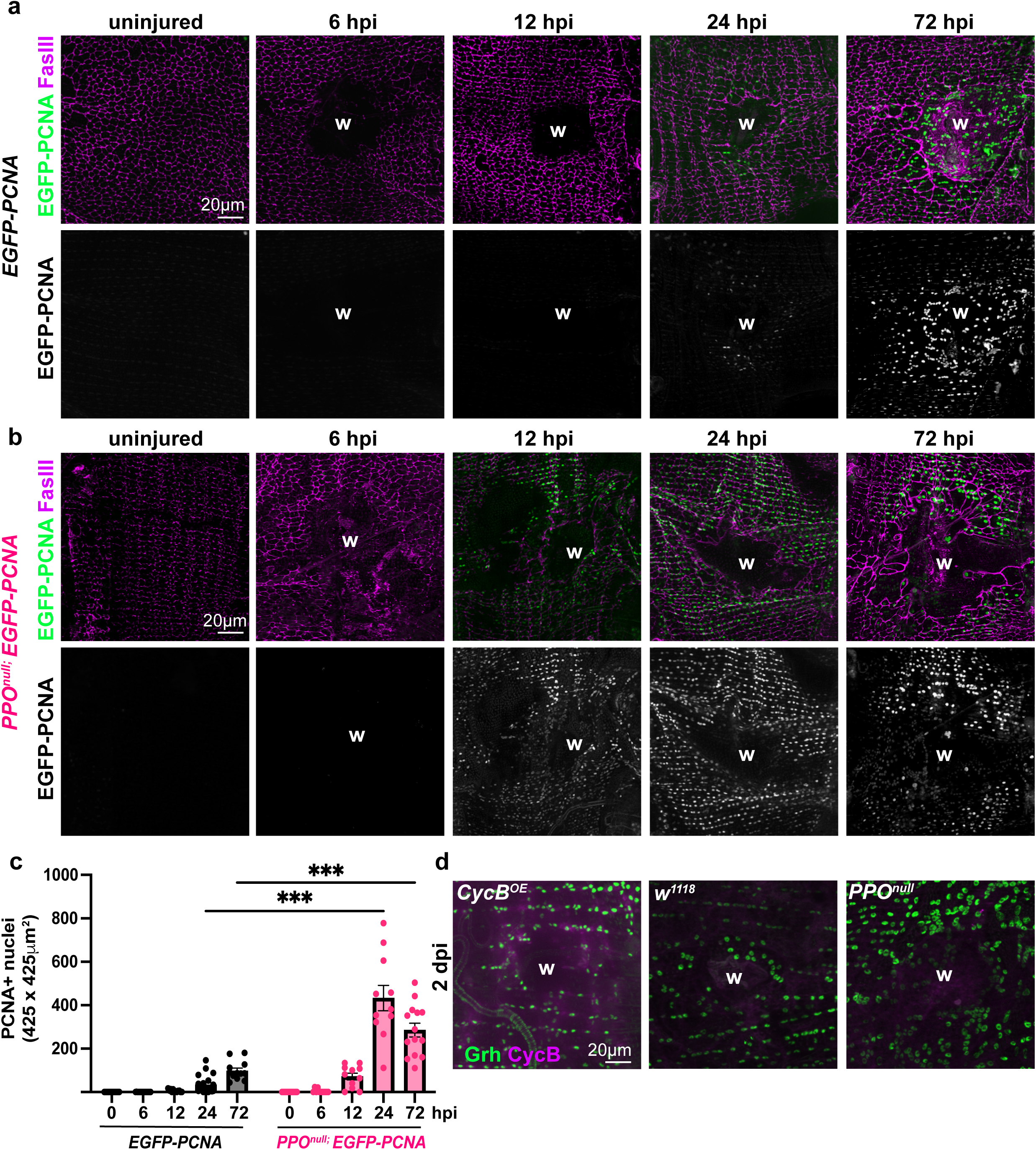
**Melanization is required to control both the extent and timing of cell cycle activity post injury**. Representative immunofluorescence images of epithelium following a time course of the S-phase marker, EGFP-PCNA in (a) *w^1118^*; *EGFP-PCNA* and (b) *PPO^null^; EGFP-PCNA* flies. Septate junctions (FasIII, magenta), EGFP-PCNA (green) and wound site (w) are indicated. (c) Quantification showing that more nuclei become PCNA^+^ post injury in the absence of PPO genes. (d) Representative immunofluorescence images of epithelium in flies overexpressing the M-phase marker, CycB (*CycB^OE^*), *w^1118^,* and *PPO^null^* flies at 2 dpi. Epithelial nuclei (Grh, green) and CycB (magenta) are indicated. Data represent the mean±s.e.m. \*\*\*\**p*<0.00001; (one-way ANOVA with Tukey’s multiple comparisons test). See Source data 4.

### The regulation of the JNK signal transduction pathway is dependent on melanization

Previous studies have demonstrated that the extent of polyploidization post injury is dependent on Jun terminal kinase (Jun) a conserved transcription factor in the JNK signal transduction pathway (Losick et al., 2016). Knockdown of Jun in the fly epithelium led to exacerbated polyploidization post injury similar to the response observed in the *PPO^null^* strain. Thus, we first assessed if JNK signaling was altered by introducing the TRE-dsRED reporter, which contains a Jun transcriptional binding site into the *PPO^null^* strain (*PPO^null^, TRE-dsRED)*. As expected, we found no expression of the TRE-dsRED reporter in uninjured wildtype or *PPO^null^* flies (Figure 5a and 5b). By 1 dpi, we found that about ∼150 nuclei around the wound expressed TRE-dsRED in controls similar to prior observations using another JNK reporter, *puc-lacZ* (Losick et al., 2013; Rämet et al., 2002). The TRE-dsRED reporter was also expressed in the abdominal lateral muscle fibers, suggesting that injury to both muscle and epithelium activates JNK signaling. However, we did not detect any TRE-dsRED expression in *PPO^null^* mutants until 2 dpi (Figure 5a and 5b), resulting in a one-day delay in JNK activation in the absence of melanization. Overall, the timing of JNK activity appears to be altered in the *PPO^null^* mutants as TRE-dsRED persisted at 4 dpi, while in the wildtype, the JNK reporter was downregulated in the epithelium, but not in the muscle cells. The expression of the TRE-dsRED reporter in the overlaying abdominal muscle fibers complicated our analysis, so we examined another JNK dependent gene, matrix metalloproteinase 1 (MMP1). Several studies in *Drosophila* have reported that the expression of MMP1 protein is dependent on JNK signaling, so we first confirmed this was the case in the adult fly epithelium as well (Huang et al., 2024; Stevens & Page-McCaw, 2012).

**Figure 5.**
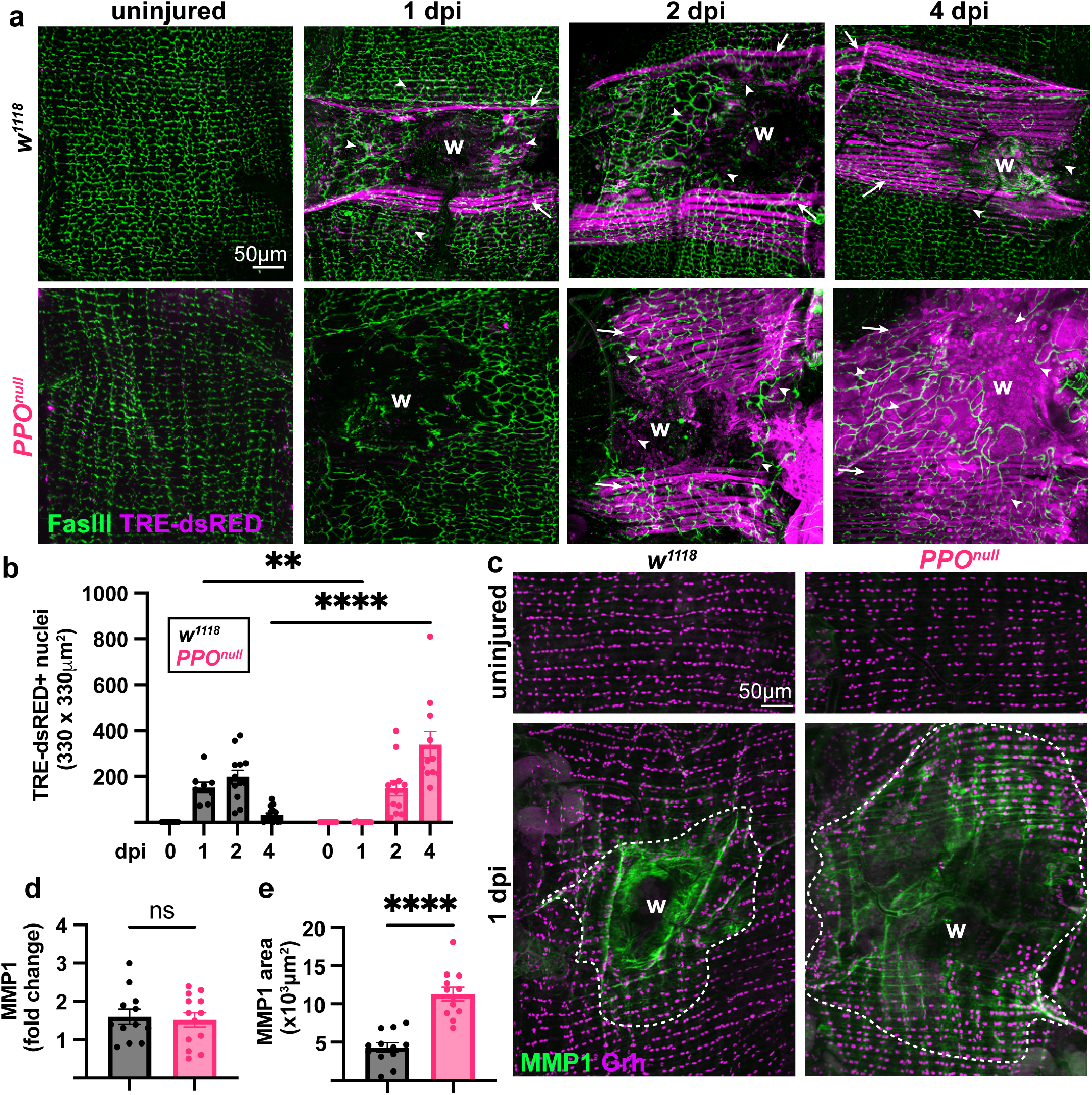
Melanization regulates the timing and expression of JNK reporters. (a) Representative immunofluorescent images of epithelium following a time course of the JNK reporter, TRE-dsRED, in *w^1118^* and *PPO^null^* fly strains. Septate junctions (FasIII, green), TRE-dsRED (magenta), and wound site (w) are indicated. Examples of TRE-dsRED+ lateral muscle fibers (arrows) and epithelial nuclei (arrowhead) are denoted. (b) Quantification of TRE-dsRED^+^ nuclei showing that TRE-dsRED expression is delayed by 1 day in the absence of PPO genes. (c) Representative immunofluorescent images of epithelium in uninjured and 1 dpi *w^1118^* and *PPO^null^* flies. Epithelial nuclei (Grh, magenta), MMP1 (green), and MMP1^+^ area (outlined, white dashed line). (d) Quantification showing fold change in MMP1 expression pre- and 1 dpi does not change significantly in the absence of PPO genes. (e) Quantification showing melanization is required to limit the MMP1^+^ area post injury. Data represent the mean ± s.e.m. ns, not significant; ****, p<0.0001 (Student’s t-test). \*\*\*\**p*<0.00001; (one-way ANOVA with Tukey’s multiple comparisons test). See Source data 5.

To do so, we overexpressed a constitutively active kinase, *Hep^CA^,* which is known to active JNK signaling in the epithelium and observed enhanced expression of both JNK reporters (Figure S2a-S2c). Likewise, we confirmed that epithelial specific expression of *Jun^RNAi^* was sufficient to reduce both TRE-dsRED and MMP1 expression post injury (Figure S2d-S2f). Despite both JNK reporters being dependent on Jun, we observed differences in their localization and temporal expression. First, MMP1 is only expressed in epithelial cells post injury and is restricted to the epithelial cells adjacent (within ∼5,000μm^2^ area) to the wound site in wildtype flies (Figure 5c and 5e). However, MMP1 protein expression was not delayed in the *PPO^null^* flies as both strains expressed MMP1 at 1 dpi (Figure 5c and 5d). The MMP1 protein extended beyond the normal range similar to TRE-dsRED reporter, suggesting that JNK signaling is indeed misregulated in *PPO^null^* mutants. The difference in temporal regulation may be due to the nature of these JNK reporters with MMP1 revealing protein expression and TRE-dsRED reporting transcriptional gene expression. Regardless, it is clear that melanization is necessary to coordinate the extent of polyploidization post injury through regulation of JNK and likely many other signal transduction pathways.

## DISCUSSION

Here, we describe a novel role for melanization in regulating wound-induced polyploidization (WIP) by limiting both cell fusion and the endocycle thus restricting cell growth post injury. Melanization also appears to be essential to restrict growth to the wound margin as polyploidization extends well beyond it in the *PPO^null^* mutants. Thus, in the absence of melanization, both the extent and magnitude of polyploidization post injury are exacerbated, leading to a defect in wound closure.

Wound healing requires coordination of acellular and cellular events, which in *Drosophila* starts with formation of a clot that remodels into a melanized scar within hours post injury (Cerenius et al. 2008; Schmid et al., 2019; Tang, 2009). The melanization at the site of injury serves multiple roles in wound healing. One as a structural support to reseal the cuticle as well as provides a scaffold for reepithelialization. While our previous studies showed that polyploidization is required for wound healing (Losick et al., 2013), here we find that polyploidization is not sufficient. The *Drosophila* adult epithelium has severe defect in closure without melanization despite the exacerbated polyploid response. In addition to serving as a scaffold, melanization is likely a signaling hub as it generates a reactive oxygen burst from the oxidation of phenols (Nappi et al., 2009; Tang, 2009). Reactive oxygen species (ROS) are known to modulate MAP kinase signal transduction pathways by reducing cysteine residues essential for kinase activity (Son et al., 2013). Therefore, an altered ROS response may contribute to the misregulation of signal transduction pathways, including JNK, as shown in this study.

Both the temporal and spatial activation of JNK signaling is altered by genetic loss of melanization. The transcriptional reporter, TRE-dsRED, is delayed which is consistent with our previous observations that Jun is required to limit polyploidization post injury. The TRE-dsRED reporter includes only the Jun transcription binding site, so it is more specific to Jun transcriptional activity than staining for MMP1 protein (Chatterjee & Bohmann, 2012). This is one plausible explanation for the difference in temporal regulation of the TRE-dsRED reporter and MMP1 immunostain in *PPO^null^* flies. The MMP1 protein is likely under the control of other transcription factors in addition to Jun. Still both JNK reporters were consistent in showing an exacerbated area of active JNK signaling further illustrating that melanization is necessary to restrict polyploidization to the wound margin. The expanded expression of MMP1 could degrade the extracellular matrix which, like in development and cellular niches, is necessary to restrain morphogens that regulate tissue growth (Page-McCaw, 2008; Waghmare & Page-McCaw, 2022).

One of the most dramatic phenotypes observed in this study was melanization’s role in regulating the timing of S-phase entry post injury. Here, we find that melanization is critical to restrict the endocycle post injury by preventing S-phase entry until 24 hpi.

We postulate that loss of the melanin scar disrupts epithelial differentiation as we found Grh expression was reduced in the epithelial nuclei within the syncytia. Grh is a conserved transcription factor required for expression of genes in epithelial differentiation in flies and mice (Gasperoni et al., 2022; Paré et al., 2012). In addition, Grh has been reported to regulate expression of PCNA, an DNA replication protein (Hayashi et al., 1999), so its misexpression may contribute to the altered endoreplication observed within and outside the central giant syncytium. Taken together, we have shown that melanization plays a key role in regulating WIP by confining epithelial competence to re-enter the cell cycle to the site of injury and coordinate signaling to facilitate wound healing.

## Data Availability Statement

The authors affirm that all data necessary for verifying the conclusions are present within the article, figures, and Supplemental Information. All raw data points reported in figures can be found in the Source data files. Additionally, the *Drosophila* strains and antibodies used in this study are listed Table S1 and S2 and are available upon request or as referenced.

## Acknowledgments

We would like to thank the members of the Losick lab, including Lydia Bischoff, Sophie Jalkut, Minqi Shen, Carmen Uribe, and Jack Weidenbach for critical review of this manuscript. The fly strains used in this study were purchased from (with support by): the Bloomington Drosophila Stock Center (NIH P40OD018537) and Vienna Drosophila Resource Center. The FasIII and MMP1 antibodies used in this study was purchased from the Developmental Studies Hybridoma Bank (created by the NICHD and maintained at The University of Iowa, Department of Biology, Iowa City, IA 52242, USA). Images were acquired using microscope supported by the National Science Foundation under Grant No. 1626072 with additional thanks to Bret Judson whom directs the Boston College Imaging Core for infrastructure and support.

## Authors contributions

Conceptualization: L.G. and V.P.L.; Methodology: L.G. and V.P.L.; Validation: L.G.; Formal analysis: L.G. and V.P.L.; Investigation: L.G., E.M., and L.M.; Resources: L.G. and V.P.L. Writing - original draft: L.G. and V.P.L.; Writing - review & editing: L.G., E.M., L.M., and V.P.L.; Visualization: L.G. and V.P.L.; Supervision: V.P.L.; Project administration: V.P.L.; Funding acquisition: V.P.L.

